# The patterns of vascular plant discoveries in China

**DOI:** 10.1101/381731

**Authors:** Muyang Lu, Lianming Gao, Hongtao Li, Fangliang He

## Abstract

Botanical discovery has a long journey of revelation that contributes unparalleled knowledge to shape our understanding about nature. Plant discovery in China is an immanent part of that journey. To understand the patterns of plant discoveries in China and detect which taxa and areas harbor most numbers of undiscovered species, we analyzed the discovery times of 31093 vascular plant species described in *Flora of China*. We found that species with larger range size and distributed in northeastern part of China have a higher discovery probability. Species distributed on the coast were discovered earlier than inland species. Trees and shrubs of seed plants have the highest discovery probability and ferns have the lowest discovery probability. Herbs hold the largest number of undiscovered species in China. Most undiscovered species are found in southwest China, where three global biodiversity hotspots locate. Spatial patterns of mean discovery year and inventory completeness are mainly driven by the total number of species and human population density in an area and whether the area is coastal or not. Our results showed that socio-economic factors dictate the discovery patterns of vascular plants in China. Undiscovered species are mostly narrow-ranged, inconspicuous endemic such as herbs, which are prone to extinctions and locate in biodiversity hotspots in southwest China. We suggest that the future effort on plant discovery in China be prioritized in southwest China.

## INTRODUCTION

Despite 270 years’ discovery of species since Linnaeus, our knowledge about the biological diversity in nature is still far from complete. According to a recent estimate, fewer than 20% of species on earth have been so far discovered (Mora *et al*., 2011). With the ongoing biodiversity crisis, this lack of knowledge has become a major obstacle to biodiversity conservation as many species could go extinct without ever being known to science (Costello, May, & Stork, 2013; Humphreys *et al*., 2019).

Early botanical and zoological discoveries are often biased towards large-sized, charismatic species with widespread geographic distributions (Gibbons *et al*., 2005; Ferretti *et al*., 2008; Stork *et al*., 2008, 2015; Essl *et al*., 2013; Randhawa, Poulin, & Krkošek, 2015). Positive correlations between species discovery probability and body size have been found in a variety of taxa including insects (Gaston & Hudson, 1994), birds (Blackburn & Gaston, 1995), mammals (Paxton, 1998; Medellín & Soberón, 1999; Collen, Purvis, & Gittleman, 2004), fishes (Zapata & Robertson, 2007), but not in taxa like reptiles (Reed & Roback, 2002) and marine holozooplanktons (Gibbons *et al*., 2005). More recently discovered species are often of greater conservation interest because they are more likely to be narrow ranged and rare, and thus are prone to extinctions (Diniz-Filho *et al*., 2005; Bebber *et al*., 2007a, 2010; Tedesco *et al*., 2014; Xu *et al*., 2019).

In addition to biological factors, species discovery is also influenced by human factors such as taxonomic effort, technology innovations, and social-economic events. For example, it is commonly observed for many taxa that species discovery rates dropped during the two World Wars and peaked in the 1990s with the emergence of molecular techniques (Gaston, 1995; Bebber *et al*., 2007b; Joppa, Roberts, & Pimm, 2011b; Lu & He, 2017). Geographically speaking, Europe and North America have the most complete species inventories due to their long histories of exploration (Gaston, 1995; Essl *et al*., 2013), while species discovery in less explored continents such as South America and Africa was affected by colonization histories and indigenous knowledge (Gaston, 1995; Diniz-Filho *et al*., 2005; Rosenberg *et al*., 2013; Ballesteros-Mejia *et al*., 2013). As a result, biodiversity hotspots, most of which locate in less developed countries (Myers *et al*., 2000), often harbor the largest number of undiscovered species (Joppa *et al*., 2011a; Giam *et al*., 2012). This imposes a more serious challenge for biodiversity conservation in developing regions where economic growth is often achieved at the expense of environmental degradation (He, 2009).

China harbors nearly one tenth of the plant species on earth (Joppa *et al*., 2011b; Lu & He, 2017). However, its rapid economic growth over the past three decades has resulted in the colossal loss of millions of hectares of natural habitats (He, 2009). The sustainable development of China depends on balancing economic growth and preservation of natural habitats. Knowing where undiscovered species may locate is necessary for making decision on habitat protection and conservation managements.

The interest for cataloging species in China has long predated the invention of Linnaeus’ binomial nomenclature, but in the light of modern taxonomy, much credit should be given to western naturalists who diligently collected specimens and described species since the first arrival of Jesuits in the 16^th^ century, as reflected by the fact that nearly 70% of the type specimens of Chinese plants are kept in herbaria in Europe and America (Chen, 1994). Due to logistic constraints and political instability, most naturalists in the 18^th^ and 19^th^ centuries made their botanical collections in the coastal areas (Bretschneider, 1898; Fan, 2004), which likely had affected the patterns of discovery records.

There are more than 31000 vascular plant species documented in China, but this inventory is not complete and many new species still await discovery (Lu & He, 2017). Knowing what are those inconspicuous species and where they may locate is important to discover them. Therefore, the objectives of this study are: (1) to model and map geographic variation in botanical discovery of vascular plant species in China, (2) to find out which taxa have the largest numbers of undiscovered species and where the undiscovered species most likely are, and (3) to quantify what factors (e.g., species’ range size and growth form) may influence the spatial distribution of plant discoveries in China. This study will contribute to understanding the pattern of species discoveries and their underlying factors and should be of significance to botanical discoveries of other regions beyond China. The identification of taxonomic and geographic gaps of undiscovered species will facilitate prioritizing our taxonomic and conservational efforts in the future.

## MATERIALS AND METHODS

### Data

Data including species names, authorities, discovery years, province-level biogeographic distributions, and genus-level growth forms were compiled from *Flora of China* (FOC) (Data available in Appendix S2) which has a totol number of ~31000 species (http://efloras.org). We treated discovery year as the time a species was first described in any name in a scientific publication. When estimating the number of undiscovered species, data after 2000 were excluded as a routine to avoid the effect of delayed entrance of newly discovered species (Costello & Wilson, 2011; Costello, Wilson, & Houlding, 2012). Human population densities on province-level were obtained from *2010 Population Census of The People’s Republic of China* (http://www.stats.gov.cn/), and *Monthly Bulletin of Interior Statistics* (http://sowf.moi.gov.tw/stat/month/elist.htm). Province areas were obtained from *National Fundamental Geographic Information System of China* (http://nfgis.nsdi.gov.cn/nfgis/). There are in total 28 provinces after merging independent municipalities such as Beijing and Shanghai to adjacent provinces. Range size, maximum/minimum latitude, maximum/minimum longitude and whether or not a species is distributed in coastal areas were obtained from province-level distributions. Genus-level growth forms were categorized as fern, herb, shrubs/trees and vine/liana. Shrubs and trees were categorized as one group because many species have both shrubs and trees as growthforms. Vines and liana include herbaceous vines, woody lianas and all other plants with climbing forms. When a genus has several different growth forms, we used the major growth form (which has the largest number of species within the genus) as the genus-level growth form. Range size, maximum/minimum latitude and maximum/minimum longitude were standardized to [0, 1] in order to calculate the effect size on discovery probability. Range size was log-transformed before standardization. Correlations between explanatory variables were checked prior to analysis. No collinearity was found among explanatory variables (the highest correlation was 0.7).

### Cox proportional hazard model

We first modeled the discovery time using survival analysis in which the discovery time was treated as a random variable. Discovery time is the time it takes to discover a species, i.e., how long the species can “survive” not to be discovered. Because only discovered species could be recorded, there are no censored data in our study. In this case, the empirical survival curve is just the inverse of the accumulation curve (Steyskal, 1965; Essl *et al*., 2013). Cox proportional hazard (Cox PH) model was used to model the instantaneous discovery probability which describes the probability that a species will be seen in the next step of time (*t*+Δ*t*) if it remains unseen up to time *t*. This probability is expressed as a hazard function, *h*(*t*), given as

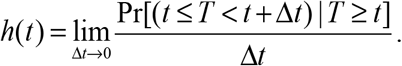

The Cox PH regression model is:

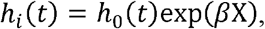

where matrix *β*X is the linear component of the model (i.e., coefficients and dependent variables). *h_i_*(*t*) is the hazard function for species *i* (i.e., the “risk” to be discovered), and *h*_0_(*t*) is the baseline hazard function. Therefore, Cox PH model is a type of generalized linear model. Compared to accelerated failure time model (AFT), Cox model is equivalent to a semi-parametric model which makes no assumption about the underlying distribution of survival time (i.e., discovery time), which is more appropriate for our data because discovery process is highly influenced by historical events (Lu & He, 2017) and the probability distribution of discovery time is unknown. Because it happened that multiple species were discovered in the same year, Efron approximation was used to break ties in discovery time (Hertz-Picciotto & Rockhill, 1997). The proportional hazard assumptions were examined by plotting the scaled Schoenfeld residuals, denoted as *β*(*t*), against time. A horizontal trend of *β*(*t*) implies that the time-independent coefficient assumption is met. Model selection was done by choosing the model with the minimum AIC value. Concordance statistic (*C* statistic) was used to show the discriminative ability of a model. It is equivalent to the area under the Receiver Operating Characteristic Curve (AUC) in logistic regression, with the value of 0.5 indicating no discrimination power and the value of 1 indicating perfect discrimination. We also presented the fitted survival curves for different treatments (i.e. inland or coastal distributed species, and species with different growth forms) using strata models (in strata model each treatment has a different baseline function *h*_0_(*t*)). Effect sizes are regression coefficients of the standardized predicting variables. All survival analyses were conducted using R package ‘survival’ (Therneau, 2015).

### Estimating species richness

Because models that take into account of taxonomic effort (Joppa *et al*., 2011a,b; Lu & He, 2017) did not converge at the province level, we used a modified logistic discovery model (Lu & He, 2017) to estimate species richness for different growth forms and for each province:

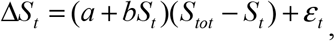

where Δ*S_t_* is the number of species discovered per time interval (5 years in this study), *S_tot_* is the total number of species in a region, *S_t_* is the accumulative number of species discovered up to time *t* (=0, 5, 10, 15, 20, … years), *a* and *b* are fitting parameters and *ε_t_* is the error term. Our goal is to estimate *S_tot_*. Note that the logistic discovery model is different from the logistic regression model. The ‘logistic’ part of the discovery model derives from the logistic shape of species discovery curve. The logistic discovery model provides only conservative estimates in certain cases (Bebber *et al*., 2007b; Essl *et al*., 2013; Lu & He, 2017). Inventory completeness was calculated as the ratio of the number of discovered species to the estimated total number of species in a province.

### Spatial patterns of discovery time and inventory completeness

We further examined the relationship between mean discovery time (i.e., the average number of years taken to discover the recorded species in an area) of a province and human population density, total number of species, whether a province is on the coast or inland, and province area using ordinary linear regression. The spatial autocorrelation of mean discovery year on the province scale was examined by Moran’s *I*. The neighborhood structure of provincial polygons is defined by contiguity (only polygons with shared borders are counted as neighbours). We proceeded with ordinary linear regression after no spatial autocorrelation was detected in the residuals of the model. Beta regression was used to model inventory completeness with the same set of covariates as modeling discovery time (i.e., population density, total number of species, whether coastal province or not, and area of each province) using R package ‘betareg’ (Cribari-Neto & Zeileis, 2010). Beta regression is preferred because it only assumes that the response variable ranges from 0 to 1 and is flexible to accommodate the shape of the distribution (symmetric or skewed). Logistic regression is for handling binary data, not for proportional data. If logistic regression is used for proportional data, the base counts that lead to the calculation of proportion must be known. Without that information, it is inappropriate to use logistic regression. In our data, the base population which is the true total number of species is not known and hence logistic regression is not applicable.

## RESULTS

### Cox proportional hazard model for species discovery probability

The results in Table 1 show that ferns have the lowest discovery probability and trees and shrubs have the highest discovery probability among all taxonomic groups. For example, in year 1953 ferns have the highest proportion of undiscovered species and trees and shrubs have the lowest proportion of undiscovered species (Fig. 1A). Coastal species were discovered earlier than inland species (Fig. 1B). The discovery probability of a species increases with its range size, maximum latitude and maximum longitude. The effect sizes of maximum latitude, minimum latitude, maximum longitude and minimum longitude are larger than the effect sizes of range size, coastal distribution and growth form, suggesting the importance of geographic locations to discovery probability (Table 1).

**Table 1.** Cox proportional hazard model with all variables (growth form as categorical data and tree/shrub used as the baseline variable).

|  | Effect size | Standard error | *P*-value |
| --- | --- | --- | --- |
| Growth form.fern | -0.82 | 0.03 | <0.001 |
| Growth form.herb | -0.13 | 0.01 | <0.001 |
| Growth form.vine.liana | -0.13 | 0.03 | <0.001 |
| Range size | 0.51 | 0.06 | <0.001 |
| Coast | 0.13 | 0.02 | <0.001 |
| Maximum longitude | 1.43 | 0.05 | <0.001 |
| Minimum longitude | -1.05 | 0.05 | <0.001 |
| Maximum latitude | 0.83 | 0.07 | <0.001 |
| Minimum latitude | -0.77 | 0.04 | <0.001 |
| N = 30944 |  |  |  |
| Concordance = 0.653 (se = 0.002) |  |  |  |

**Figure 1.**
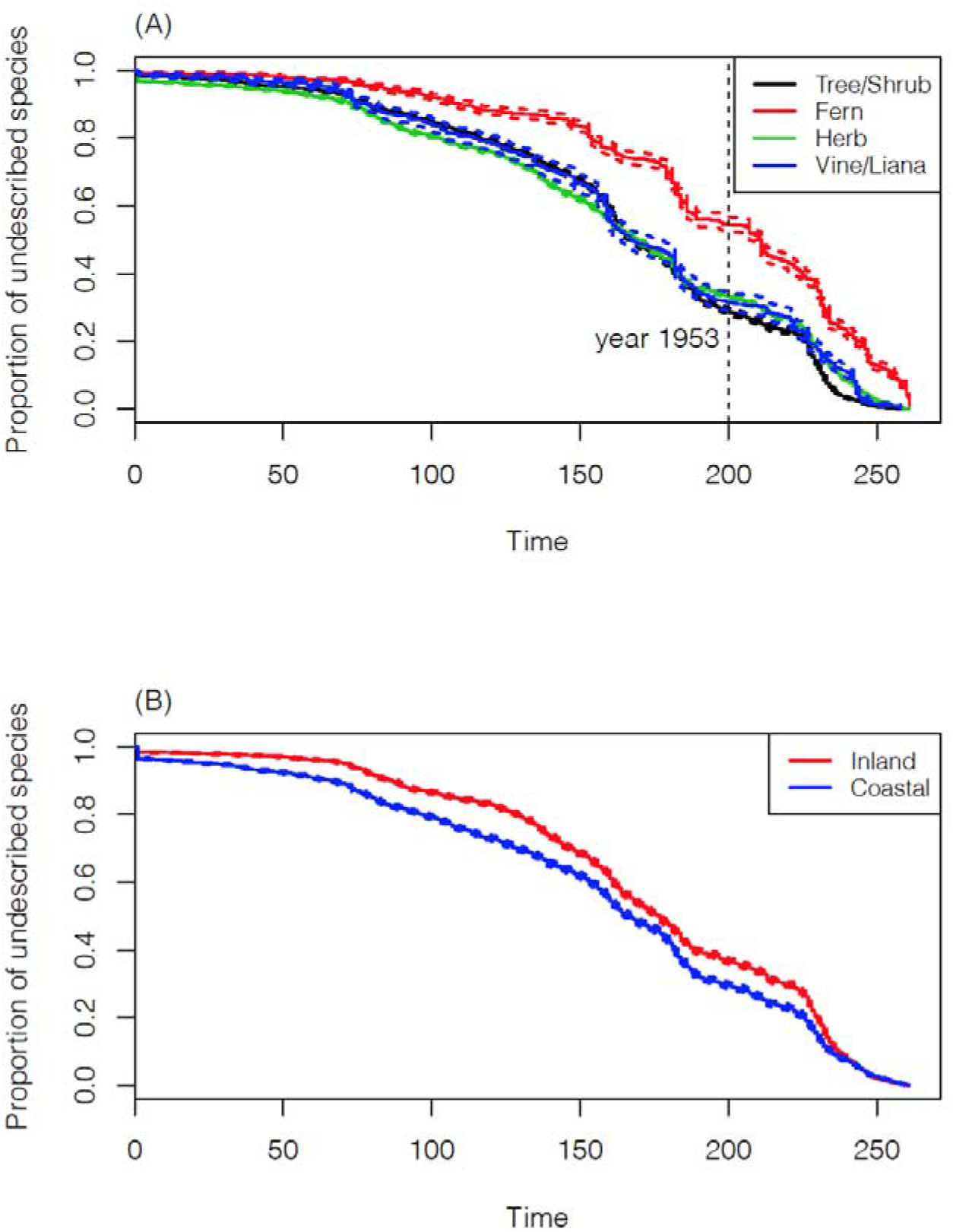
Fitted survival curves of the Cox proportional hazard models (A) stratified on variable ‘growth form’ and (B) stratified on variable ‘coast’. Dashed lines show 95% confidence intervals.

### Estimating species richness

The accumulation curve (i.e. the inverse of survival curve) of fern species has the steepest slope at recent years (Fig. 2A), suggesting that a considerable number of fern species remain to be discovered in the future. The logistic model estimates that the inventory completeness of ferns is 0.62. For seed plants, the estimated inventory completeness is 0.73 for herbs, 0.75 for shrubs and trees, and 0.68 for vines/lianas (Table 2). Herbs harbor the largest number of undiscovered species. The low ferns discovery rate is reflected by the fact that it took on average the longest time to discovery a fern species, while herbs were earlier to be discovered (*p* < 0.05 for all pairs except between tree/shrub and vine; Fig. 2E).

**Figure 2.**
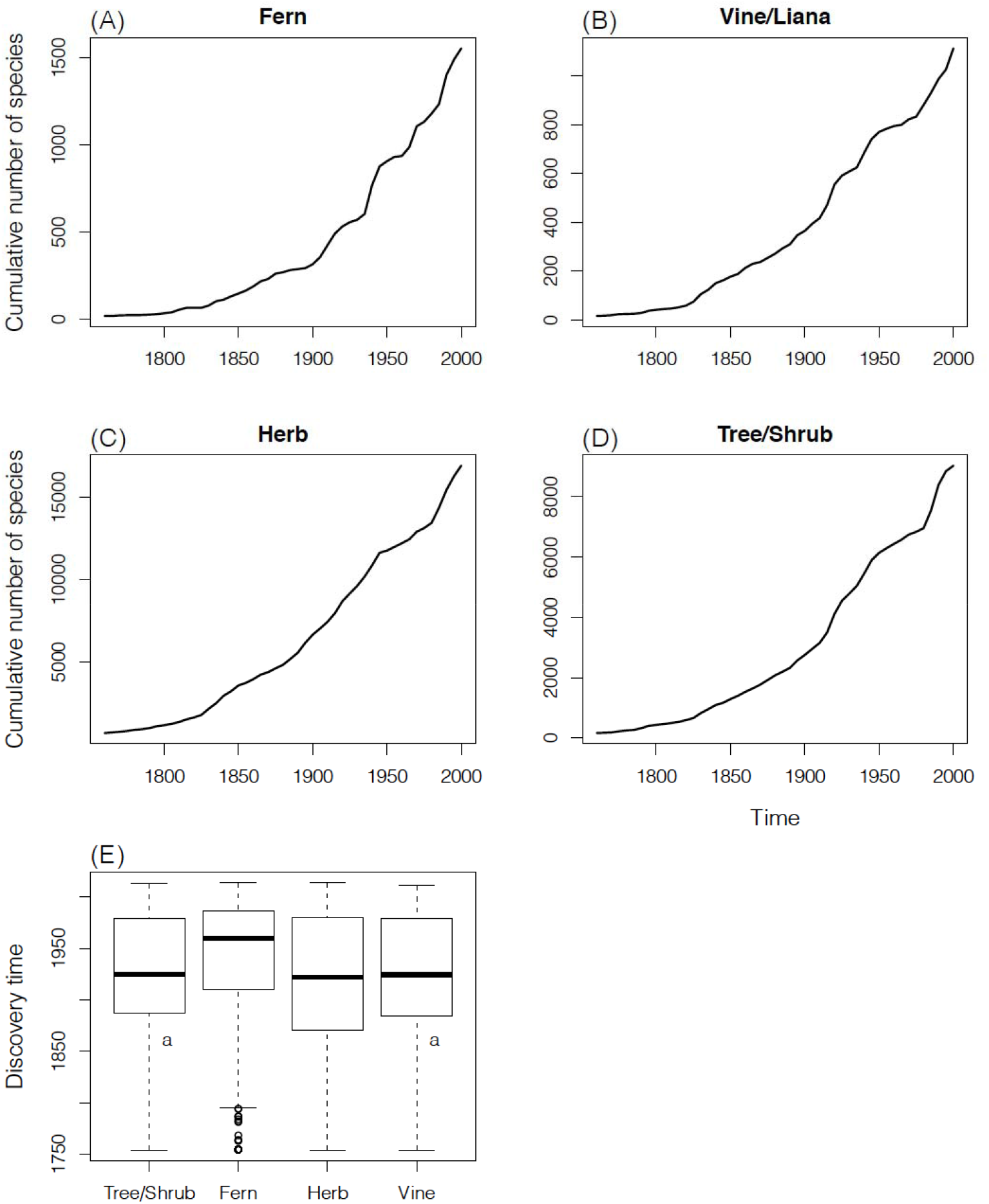
(A) – (D) Species accumulation curves for four different taxonomic groups (based on 5 years-interval data). (E) Boxplot for discovery time of the four growth forms. “a” labels the groups with no significant difference in the TurkeyHSD test.

**Table 2.** Estimated species richness for different growth forms.

|  | Number of<br>discovered<br>species | Estimated<br>total number<br>of species | Lower 95%<br>bound | Upper 95%<br>bound | Completeness<br>(lower 95% - ) |
| --- | --- | --- | --- | --- | --- |
| Fern | 1921 | 2712 | 749 | 4676 | 0.622 (0.41-) |
| Vine/Liana | 1202 | 1710 | 856 | 2563 | 0.685 (0.47-) |
| Herb | 18370 | 23867 | 16982 | 30752 | 0.737 (0.59-) |
| Tree/Shrub | 9451 | 12998 | 8855 | 15741 | 0.754 (0.60-) |

### Spatial patterns of mean discovery year and inventory completeness

The mean discovery year increases from northeast towards southwest (Fig. 3A). Besides geographic variables (latitudes and coastal distribution), the best model shows that the total number of species in a region also has a significant effect on mean discovery year (Table 3). The total number of species is positively correlated with mean discovery year (Fig. 4B). Mean discovery year decreases with latitude (Fig. 4C) and is earlier in coastal provinces than in inland provinces (Fig. 4D).

**Figure 3.**
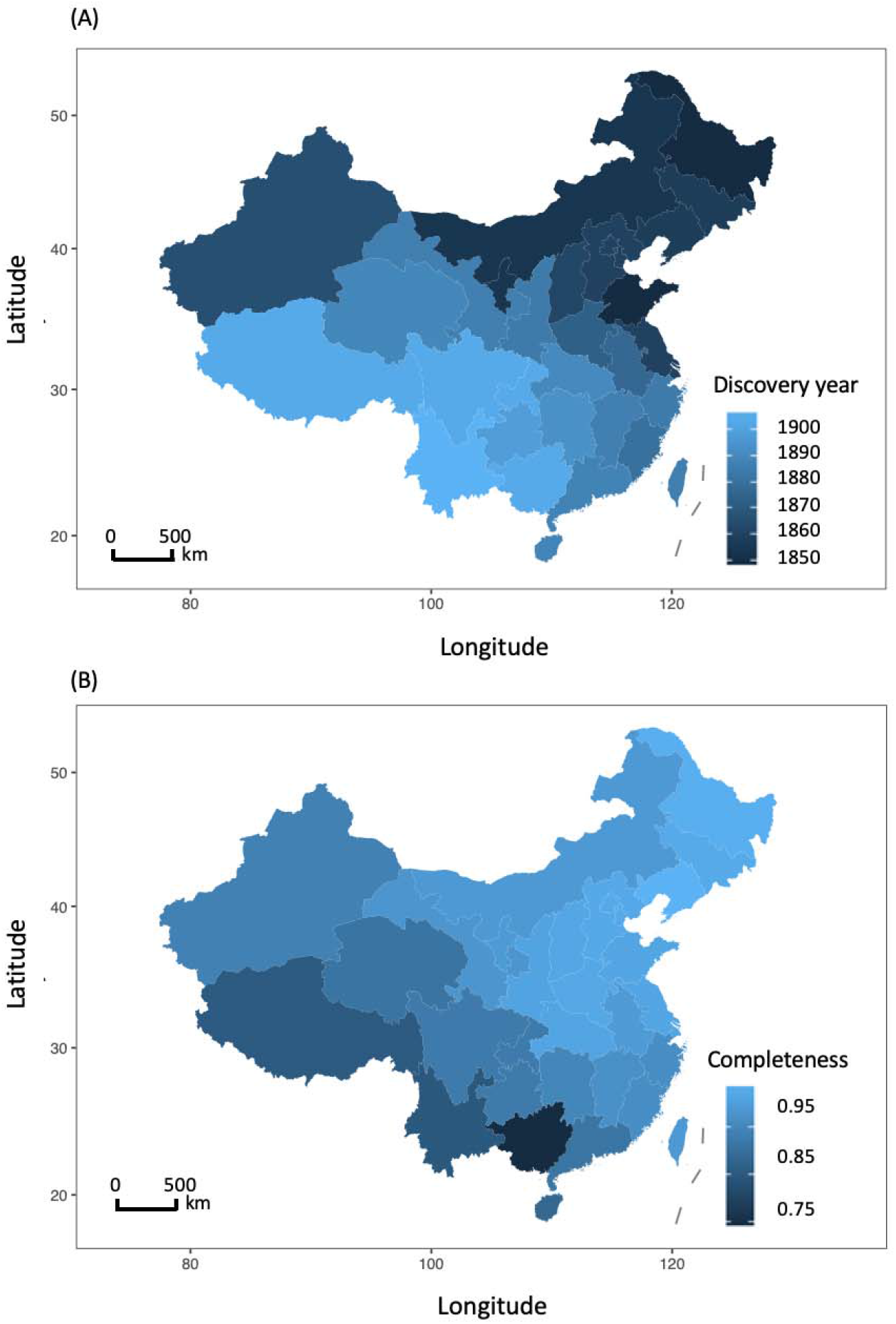
(A) Distribution of mean province-level species discovery year in China. (B) Distribution of species inventory completeness. Mollweide projection is used for the distributions.

**Table 3.** Linear regression of mean discovery year.

|  | Coefficient | Standard Error | *P*-value |
| --- | --- | --- | --- |
| Intercept | 1941 | 1.5 | <0.001 |
| Number of species | 0.0029 | 0.0008 | <0.01 |
| Coast | -13.8 | 4.49 | <0.01 |
| Mean latitude | -2.03 | 0.36 | <0.001 |
| Multiple $R^2 = 0.82$, Adjusted $R^2 = 0.80$ | | | |

**Figure 4.**
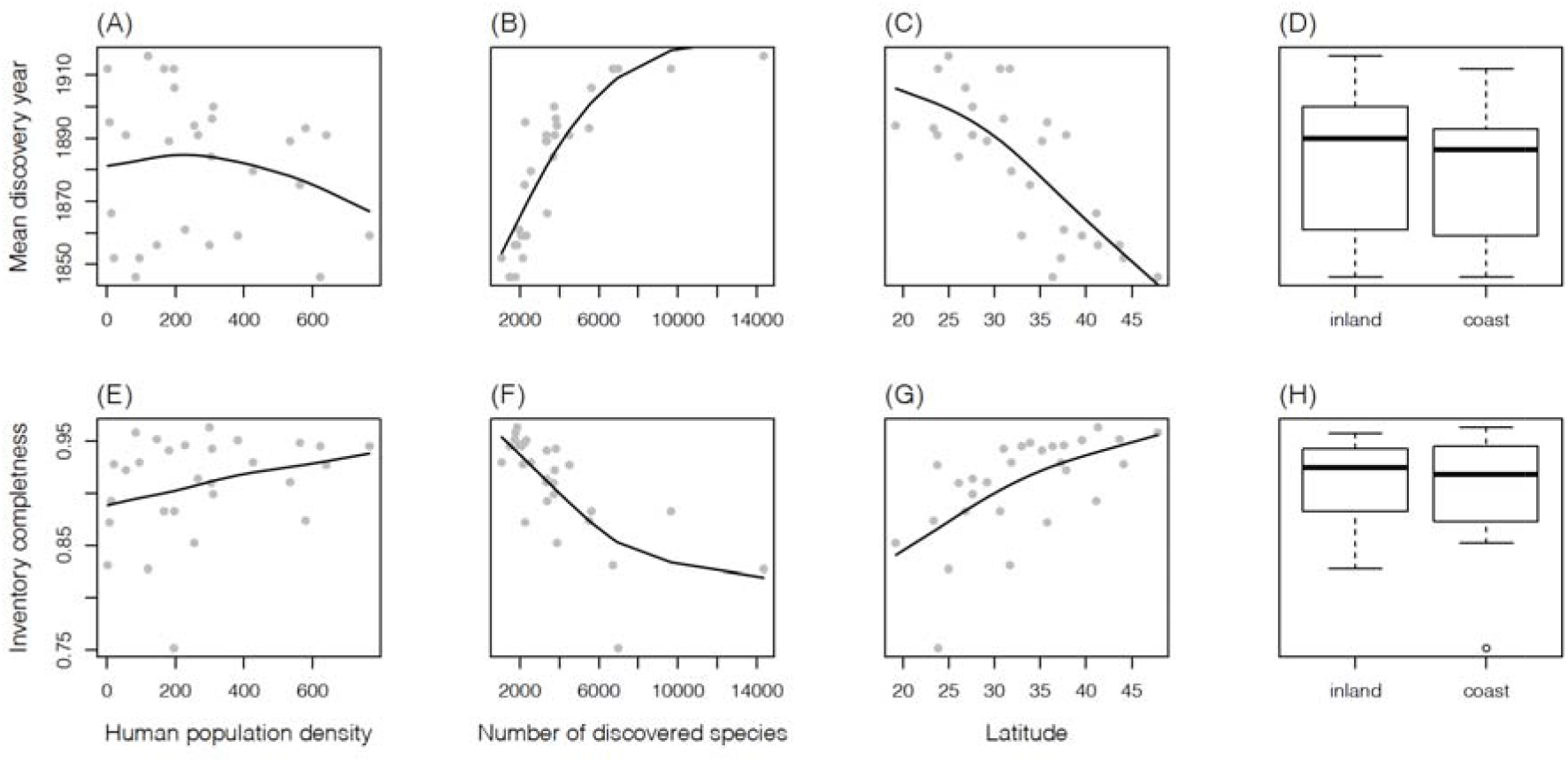
(A) – (C) Mean species discovery year against human population density, number of discovered species, and latitude. (D) Boxplot of mean discovery year for inland and coastal provinces. (E) – (G) Species inventory completeness against human population density, number of discovered species, and latitude. (H) Boxplot of inventory completeness for inland and coastal provinces. Solid curves show the fitted smooth spline curves.

We estimate that in 18 of the 28 provinces, plant species discoveries are more than 90% complete. Provinces with the largest proportion of undiscovered species are in the southwest China (75.1% in Guangxi province and 82.7% in Yunnan province; Fig. 3B). Beta regression shows that inventory completeness is positively correlated with human population density, and negatively correlated with area and latitude of a province (Table 4).

**Table 4.** Beta regression of province level inventory completeness.

|  | Coefficient | Standard Error | *P*-value |
| --- | --- | --- | --- |
| Intercept | -0.39 | 0.23 | <0.05 |
| Human population density | 8.70e-04 | 3.17e-04 | <0.001 |
| Area | -5.23e-07 | 1.33e-07 | <0.001 |
| Latitude | 0.07 | 6.8e-03 | <0.001 |
| Pseudo $R^2 = 0.85$ | | | |

## DISCUSSION

Today’s knowledge about biodiversity is the result of arduous quest of generations of naturalists for botanical and zoological discoveries. Although the nomenclatures of species are universally binomial, the stories behind their discoveries are not and many of them are as colorful as the species that were discovered (Kilpatrick, 2014). It is however surprising that the literature in ecology is largely disconnected from this rich history of species discoveries (but see Collen *et al*., 2004; Diniz-Filho *et al*., 2005; Randhawa *et al*., 2015). Worse still is that the data of species discoveries are not much used for answering questions important to biogeography and biodiversity studies although there have been efforts in using such data for estimating regional and global species richness (Gaston, 1995; Joppa *et al*. 2011a, b; Bebber *et al*. 2007a, b, 2014; Essle *et al*. 2013; Lu, & He 2017). The rich information provided by discovery history is especially valuable for filling the knowledge gap in biodiversity research (Meyer *et al*., 2015; Meyer, Weigelt, & Kreft, 2016) because it provides guidance about when and where future discoveries are going to be made and what traits are important to future discovery (Collen *et al*., 2004; Diniz-Filho *et al*., 2005). Therefore, knowledge on species discovery is of great value for species conversation if we strive to describe all species before their extinction (Costello *et al*., 2013; Tedesco *et al*., 2014; Humphreys *et al*, 2019). In this study, we compiled data on vascular plant species discovered over 250 years in China for understanding the geographic variation of discovery rates and for assessing the completeness of botanical inventory of the country.

Our analyses showed that vascular plant species with a larger range size is more likely to be discovered in China, consistent with the previous findings that widespread species are discovered earlier in the history (Bebber *et al*., 2007a; Essl *et al*., 2013). Tree and shrub species of seed plants have the highest discovery probability while fern species have the lowest discovery probability. Herb and vine/liana species of seed plants all have similar discovery probabilities. The unique history of botanical discovery in China is revealed by the fact that species distributed on the coast have a higher discovery probability than inland species even when geographic information such as the maximum and minimum latitude range of a species is included in the model (Table 1), and is also shown by the result that province-level mean discovery time is negatively correlated with coastal distribution (Table 3). This is likely because coastal areas in China were most economically developed and much more accessible to western naturalists since the Opium War (Bretschneider, 1898; Fan, 2004).

Our results indicate that fern is likely to be the most under-discovered taxon in China because the species discovery curve for fern shows little sign of level-off (Fig. 2A, Table 2). Herbs have the largest number of undiscovered species (Table 2) and the second lowest discovery probability estimated from the Cox proportional hazard model (Table 1). Higher discovery probability usually leads to higher inventory completeness, which is shown by the concordance between ranks of discovery probability and ranks of inventory completeness among taxa in our results (Table 1, Table 2). Although the inventory completeness estimated from this study varies among the growth forms, the difference in inventory completeness is relatively small in magnitude (~ 2% between herb and tree/shrub). Herbs have the largest number of undiscovered species likely because the total number of herb species is larger than that of any other growth form of seed plants. We suspect that the effect of growth form on discovery probability and inventory completeness at least partly reflects the difference in the availability of taxonomic expertise, especially for ferns. The description of fern species started relatively late in China (~1920s, Chen, 1994) compared to other groups likely because of the difficulty in distinguishing subtle morphological characters, different species concept compared to seed plants and the lack of taxonomic expertise at that time. Given that herbs also contain the largest number of undiscovered species and that most specimens of undescribed species have already been preserved in herbaria (Bebber *et al*., 2010), our study suggests that the limiting factor of discovering new species in China may be the lack of taxonomic expertise, which resonates with the call to address the challenge of “taxonomic impediments” (Ebach, Valdecasas, & Wheeler, 2011; Bebber *et al*., 2014).

The spatial patterns of mean species discovery time and inventory completeness are largely driven by human population density and species diversity in an area. Although human history did affect the spatial patterns of species discovery (Diniz-Filho *et al*., 2005; Rosenberg *et al*., 2013; Ballesteros-Mejia *et al*., 2013), geographic sampling bias does not change the prioritization of the current conservation efforts because the total number of species and number of discovered species are highly correlated (Joppa *et al*., 2011a; Giam *et al*., 2012; Fig. S3). We expect that new discoveries in the future will be most likely made in interior southwestern provinces with high species richness such as Xizang, Guangxi and Yunnan.

It is worth noting that the spatial pattern of species inventory completeness at the province level is opposite to the spatial pattern at the county level which shows that eastern part of China has lower inventory completeness (Yang, Ma, & Kreft, 2013, 2014). Yang *et al*. measured county-level inventory completeness using the slope of sample-based accumulation curves. Their assessment of inventory completeness might be biased by a sampling strategy that aims at collecting as many novel species as possible for a given amount of samples (Chen, 1994). We also argue that the slope of species accumulation curve is not a genuine measure of inventory completeness. Rather, it measures the variation in species composition in the samples used to construct the species-accumulation curve. Another possible reason for this discrepancy is that inventory completeness is scale-dependent. In a hypothetical scenario, even if the inventory completeness at the county level is on average 90%, the inventory completeness at the province level could be lower than 90% if most of the recorded species at the county level are common species. While Yang *et al*. (2014) argued that more efforts should be devoted to increasing botanical collections in eastern densely populated areas where county-level inventories are less complete, our study here does not support this advocacy. Instead, we suggest that future botanical collection efforts should be more allotted to the provinces of southwest China where there is high species diversity and the botanical inventory is least complete. Recent findings support our conlusion by showing that the majority of newly discovered species in China after the completion of *Flora of China* in 2013 came from the global biodiversity hotspots of Indo-Burma and Mountains of Southwest China (Cai *et al*., 2019), and southwest China (Yunnan, Guangxi, Sichuan, Xizang) and Taiwan are places where most new species were discovered during 2000 - 2019 (Du *et al*., 2020).

There are two major limitations in this study. The first one is that species discovery year is not the time when the species was first collected in the field but is the time when the species was first described in a scientific publication. This could lead to inflating discovery probability for widespread species if their type specimens were collected outside of China. To address this problem, we reran our Cox proportional hazard model using species only endemic to China (12917 species). The results show that the key drivers of discovery probability are mostly consistent but with some noticeable differences (Table S1): the effect size of coastal distribution became negative (Table S1) because the majority of these endemic species were discovered after 1900 (Fig. S1A), which removed the effect of early discoveries made by western naturalists. For the province-level analysis, excluding non-endemic species in China does not significantly change the spatial pattern of mean discovery year (Fig. S2A). Human population density became negatively correlated with the mean discovery year of endemic species; area became positively correlated with mean discovery year; the effect of coastal distribution and number of species are no longer significant (Table S2). For province-level inventory completeness, the effects of predicting variables are mostly consistent with the model that includes all species except that coastial distribution and longitude also became significant variables (Table S3).

The second limitation is that our data do not distinguish the discovery of a new species from a species renamed from a known synonym. As data including a full list of all synonyms at each time step are not available, we are not able to model the transition rate from synonyms to valid names (Alroy, 2002).

In summary, our study shows that most underdiscovered vascular plant species in China are ferns and herbs of seed plants, which are mostly narrowly distributed endemic species in the southwest biodiversity hotspots of China. Given the “taxonomic impediments” we are facing, more resources should be channeled to the training and recruitments of taxonomic expertise in these two particular groups. There is an urgency of cataloging undiscovered species in southwest mountainous areas for future conservation designs and botanical study.

## ACKNOWLEDGEMENTS

This work was funded by Sun Yat-sen University, East China Normal University, the Natural Sciences and Engineering Research Council of Canada (NSERC), the Strategic Priority Research Program of Chinese Academy of Sciences (XDB31000000) and the Science and Technology Basic Resources Investigation Program of China (Grant No. 2019FY100900). We thank Xingli Giam and an anonymous reviewer for the critical comments that greatly improved this study.

## Author contributions

FH and ML conceived the study; ML collected and analyzed the data; LG and HL provided the province-level checklist of *Flora of China;* ML led the writing with input from all authors.

## Appendices

**Table S1.** Cox proportional hazard model with all variables for endemic species in China (growth forms were coded as a categorical variable in which tree/shrub was used as the baseline variable).

|  | Effect size | Standard error | *P*-value |
| --- | --- | --- | --- |
| Growth form.fern | -0.83 | 0.04 | <0.001 |
| Growth form.herb | -0.082 | 0.02 | <0.001 |
| Growth form.vine | -0.23 | 0.10 | <0.001 |
| Range size | 1.21 | 0.03 | <0.001 |
| Coast | -0.12 | 0.03 | <0.001 |
| Maximum longitude | 1.30 | 0.10 | <0.001 |
| Minimum longitude | 0.09 | 0.09 | 0.32 |
| Maximum latitude | 0.30 | 0.11 | <0.01 |
| Minimum latitude | -0.70 | 0.08 | <0.001 |
| N = 12917 |  |  |  |
| Concordance = 0634 (se = 0.003) |  |  |  |

**Table S2.** Linear regression of mean discovery time for endemic species.

|  | Coefficient | Standard Error | *P*-value |
| --- | --- | --- | --- |
| Intercept | 1964 | 6.8 | <0.001 |
| Human population density | -0.013 | 0.0071 | 0.074 |
| Area | 2.30e-05 | 3.91e-06 | <0.001 |
| Mean latitude | -1.46 | 0.18 | <0.001 |
| Multiple $R^2$: 0.80, Adjusted $R^2$: 0.77 | | | |

**Table S3.**
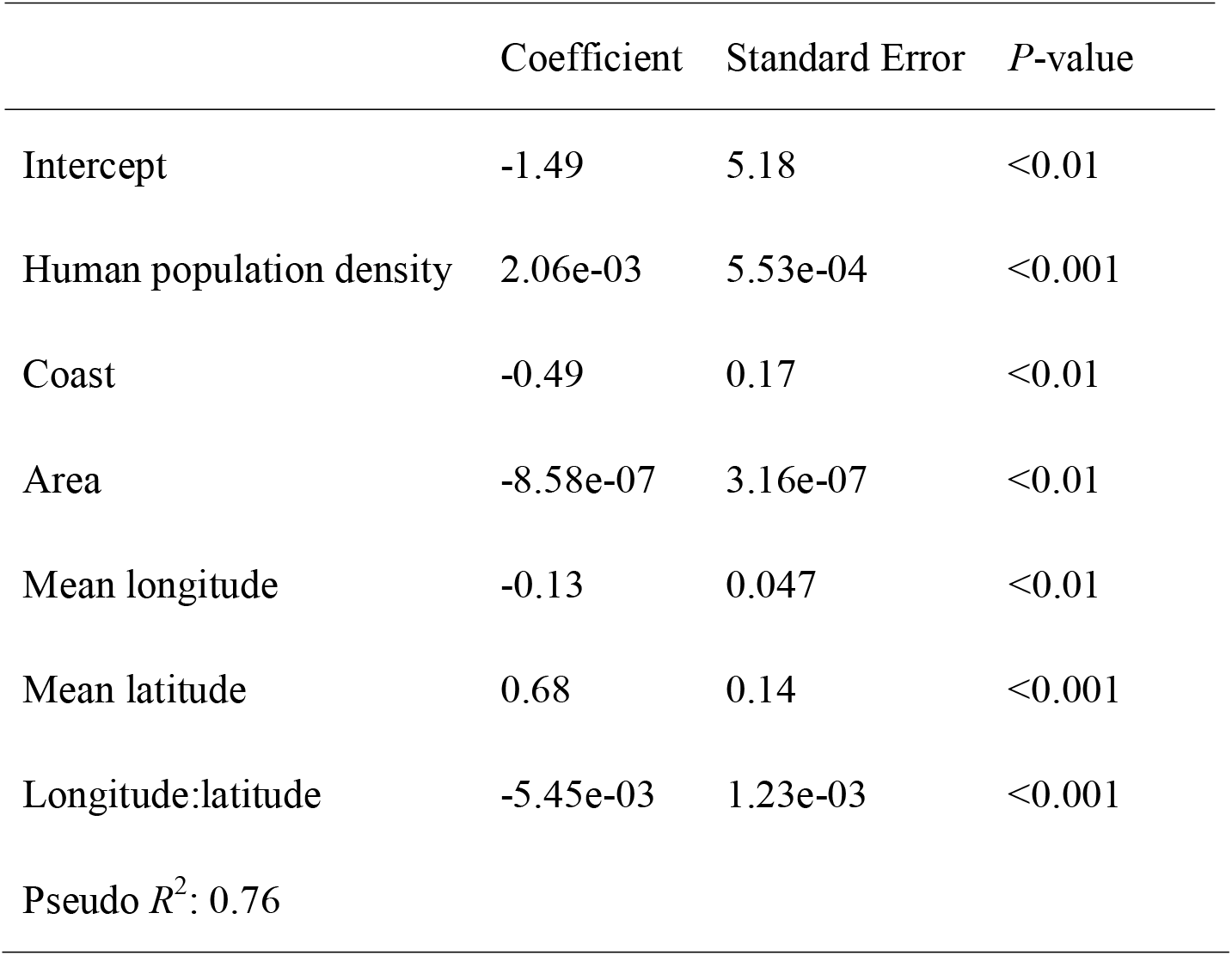
Beta regression of province-level inventory completeness for endemic species.

|  | Coefficient | Standard Error | *P*-value |
| --- | --- | --- | --- |
| Intercept | -1.49 | 5.18 | <0.01 |
| Human population density | 2.06e-03 | 5.53e-04 | <0.001 |
| Coast | -0.49 | 0.17 | <0.01 |
| Area | -8.58e-07 | 3.16e-07 | <0.01 |
| Mean longitude | -0.13 | 0.047 | <0.01 |
| Mean latitude | 0.68 | 0.14 | <0.001 |
| Longitude:latitude | -5.45e-03 | 1.23e-03 | <0.001 |
| Pseudo $R^2$: 0.76 | | | |

**Figure S1.**
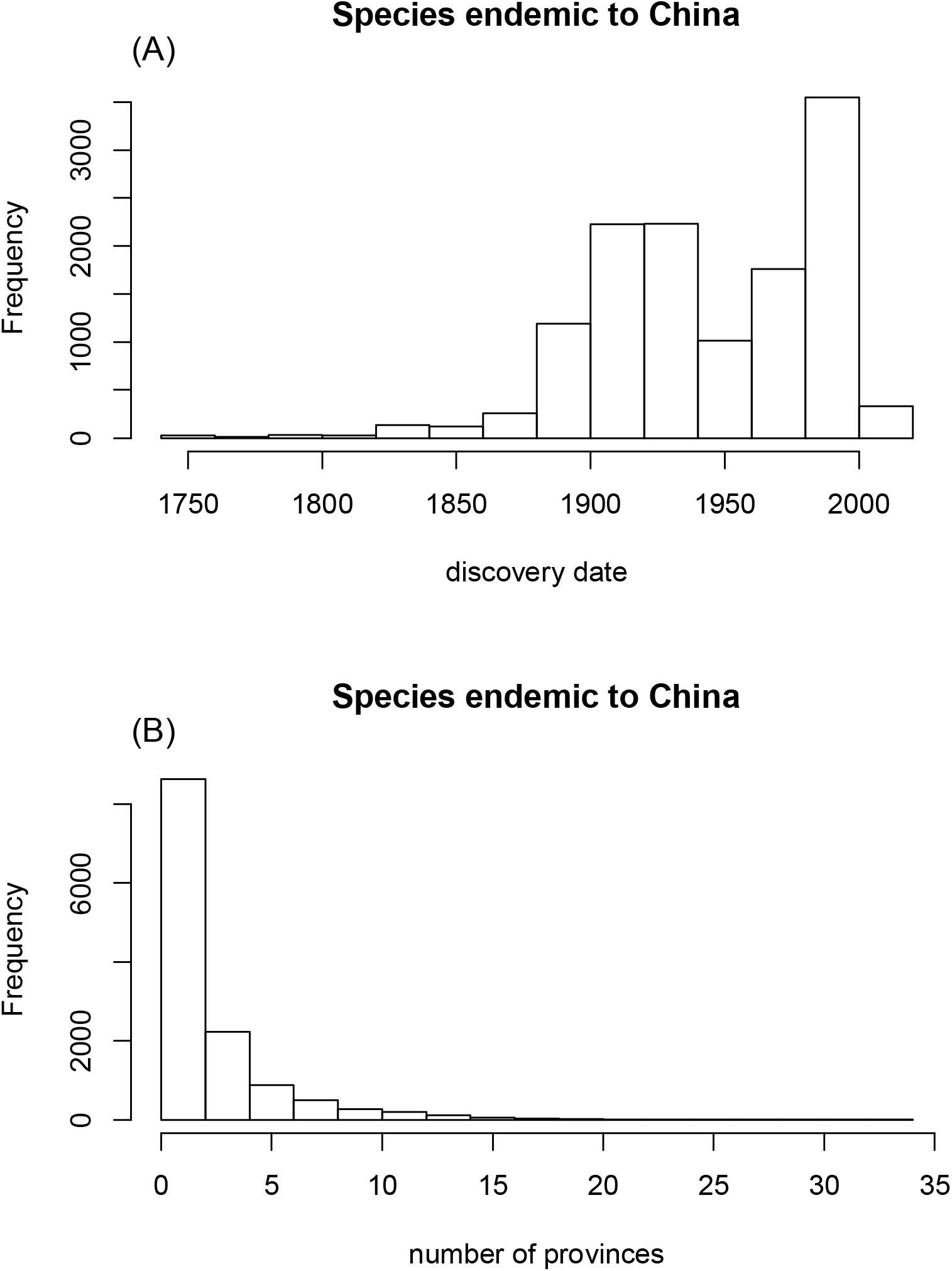
Histograms of discovery date (A) and the number of distributed provinces of endemic species to China (B).

**Figure S2.**
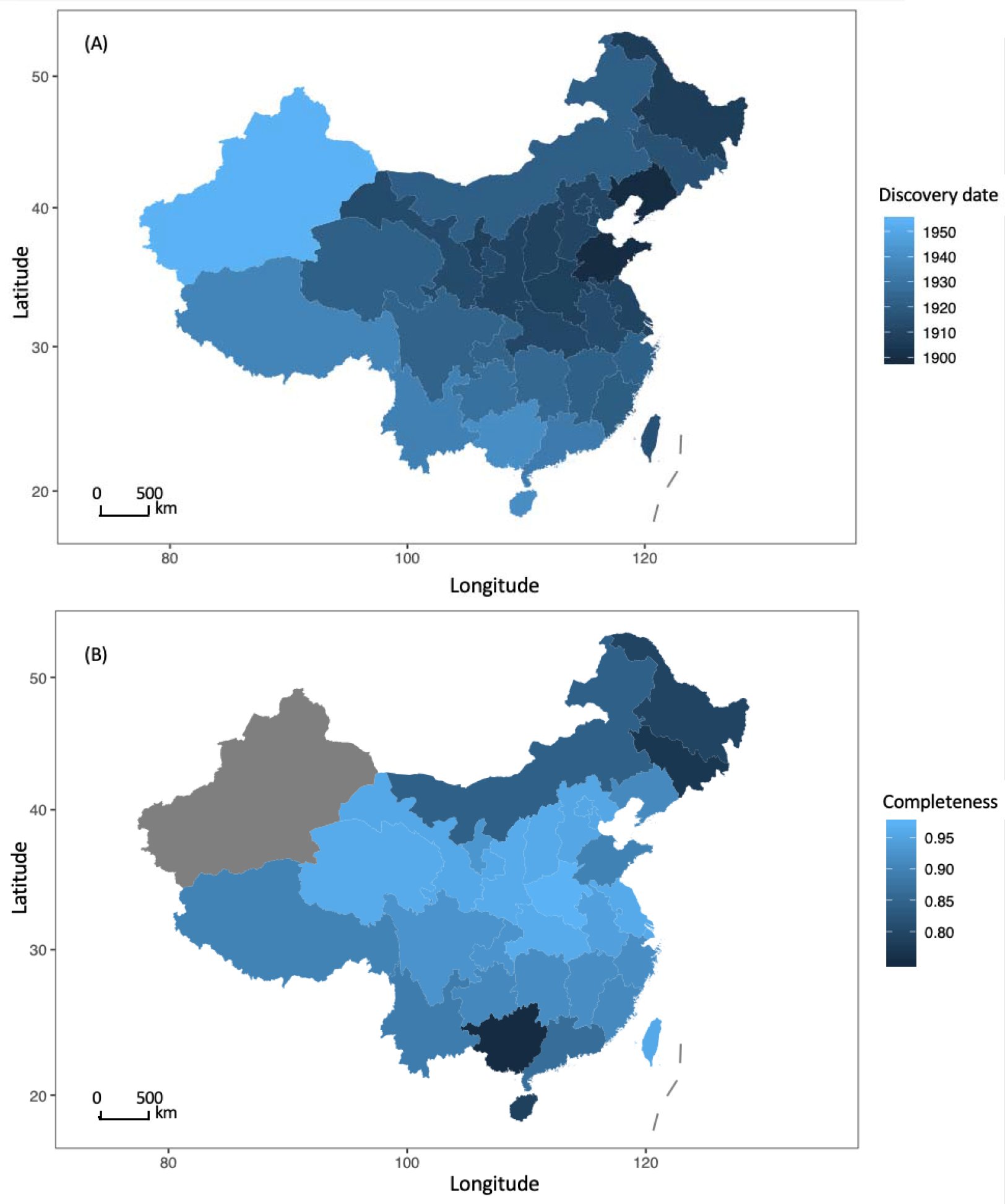
(A) Mean discovery time for province-level endemic species in China. (B) Species inventory completeness for province-level endemic species. Mollweide projection is used.

**Figure S3.**
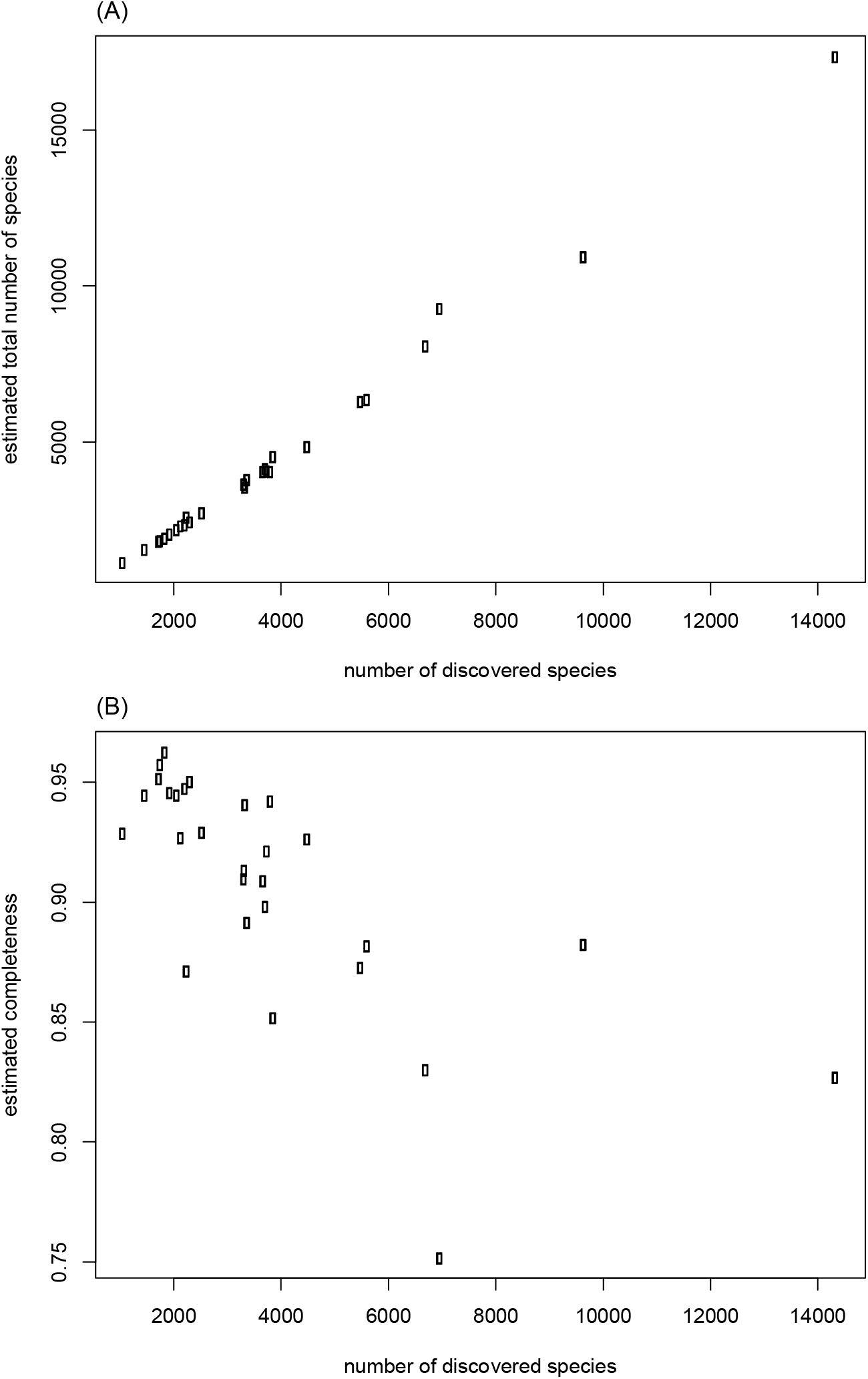
(A) Estimated total number of species against the number of discovered species at the province-level. (B) Estimated inventory completeness against the number of discovered species at the province-level.

## Notes

### Competing Interest Statement

The authors have declared no competing interest.

### Summary of Updates

Complete dataset of Flora of China is used to replace the original outdated dataset.

